# Papillomavirus can be transmitted through the blood and produce infections in blood recipients: Evidence from two animal models

**DOI:** 10.1101/541474

**Authors:** Nancy M. Cladel, Pengfei Jiang, Jingwei J. Li, Xuwen Peng, Timothy K. Cooper, Vladimir Majerciak, Karla K. Balogh, Thomas J. Meyer, Sarah A. Brendle, Lynn R. Budgeon, Debra A. Shearer, Regina Munden, Maggie Cam, Raghavan Vallur, Neil D.Christensen, Zhi-Ming Zheng, Jiafen Hu

**Affiliations:** The Jake Gittlen Laboratories for Cancer Research, Pennsylvania State University College of Medicine, Hershey, PA 17033, USA; Department of Pathology, Pennsylvania State University College of Medicine, Hershey, PA 17033, USA; Tumor Virus RNA Biology Section, RNA Biology Laboratory, National Cancer Institute, NIH, Frederick, MD 21702, USA; Department of Immunology and Microbiology, School of Basic Medical Sciences, Wenzhou Medical University, Wenzhou 325035, Zhejiang, P. R. China; Department of Comparative Medicine, Pennsylvania State University College of Medicine, Hershey, PA 17033, USA; Integrated Research Facility at Fort Detrick, National Institute of Allergy and Infectious Diseases, NIH, Fort Detrick, Frederick, MD, 21702, USA; CCR Collaborative Bioinformatics Resource (CCBR), Center for Cancer Research, NCI, NIH, Bethesda, MD 20814, USA; Advanced Biomedical Computational Science, Frederick National Laboratory for Cancer Research, Frederick, MD 21702, USA; Department of Microbiology and Immunology, Pennsylvania State University College of Medicine, Hershey, PA 17033, USA

**Keywords:** Papillomavirus, blood transfusion, animal models, CRPV/rabbit model, MmuPV1/mouse model, viral infection

## Abstract

Human papillomavirus (HPV) infections are commonly thought to be strictly sexually transmitted. However, studies have demonstrated the presence of HPV in cancers of many non-sexual internal organs, raising the question as to how the viruses gain access to these sites. A possible connection between blood transfusion and HPV-associated disease has not received much attention. We show, in two animal models, that blood infected with papillomavirus yields infections at permissive sites. Furthermore, we demonstrate that blood from actively infected mice can transmit the infection to naïve animals. Finally, we report papillomavirus infections in the stomach tissues of animals infected via the blood. Stomach tissues are not known to be permissive for papillomavirus infection, although the literature suggests that HPVs may be associated with a subset of gastric cancers. These results indicate that the human blood supply, which is not screened for papillomaviruses, could be a potential source of HPV infection and subsequent cancers.

**SUMMARY:** Human papillomaviruses cause 5% of human cancers. Currently, blood banks do not screen for these viruses. We demonstrate that blood transfused from papillomavirus-infected animals produces infections in recipients. Public health implications are significant if the same is true for humans.

**Definitions:** **Local papillomavirus infection:** An infection initiated by the direct application of virus or viral DNA to the site of infection

**Intravenous (IV) papillomavirus infection:** An infection resulting from blood-borne delivery of virus or viral DNA to the site of infection.

## INTRODUCTION

This study grew out of an observation made in 2005 (Bodaghi et al., 2005a) that a subset of children with HIV also had detectable levels of human papillomavirus (HPV) in their peripheral blood mononuclear cells (PBMCs). Some of these children were hemophiliacs who had contracted HIV through contaminated blood. All were reported to be sexually naïve. Importantly, HPV was detected in three out of the 19 seemingly healthy blood donors. A later study in 2009 demonstrated that HPV DNA is present in about 8.3% of healthy donor PBMCs in Australia (Chen et al., 2009). In addition, HPV has been detected in many malignant tissues, including the head and neck (Taberna et al., 2017), esophagus (Agalliu et al., 2018), lung (Shikova et al., 2017), colorectum (Bodaghi et al., 2005b) (Baandrup et al., 2014), prostate (Glenn et al., 2017; Tachezy et al., 2012), breast (ElAmrani et al., 2018; Malhone et al., 2018) and stomach (Mirzaei et al., 2018; Zeng et al., 2016). We asked the following questions: 1) Could blood be a non-sexual mode for the transmission of papillomavirus infections? 2) As the blood bank does not currently screen for the presence of HPV, is the public being put at risk by this omission?

HPV is strictly species-specific (Martinez and Troconis, 2014). Thus, it is not possible to study HPV infections directly in any animals. However, our laboratory is fortunate to have two preclinical animal models with their own naturally occurring papillomaviruses (Christensen et al., 2017; Doorbar, 2016; Hu et al., 2017; Uberoi and Lambert, 2017). We have developed methods that allow us to use these models to test the possibility of papillomavirus transmission by blood.

The Cottontail Rabbit Papillomavirus (CRPV) infection model has been in use in our laboratory for more than three decades. CRPV infections produce cutaneous tumors, which, in time, progress to cancer (Christensen et al., 2017). Our first series of studies for the current project was completed with this model: 1) We detected viral DNA in the blood of CRPV infected animals; 2) We infected domestic rabbits with either infectious virions or viral DNA, via the marginal ear vein, and observed tumor growth at pre-wounded back sites; 3) We drew blood from two animals that had received CRPV virions intravenously and then transfused that blood into two naïve animals via the marginal ear vein. Within ten weeks, a lesion appeared at the back of one of two tested recipients. These results demonstrate that blood containing active papillomavirus could transmit the virus to the wounded epithelium of a naïve host via the circulatory system. *These early studies confirmed the possibility that papillomaviruses can indeed be transmitted by blood and give rise to infections at receptive sites in naïve hosts.*

Next, we extended these studies to the mouse papillomavirus (MmuPV1) model, which has been under development over the past seven years in our and other laboratories (Hu et al., 2017; Uberoi and Lambert, 2017). MmuPV1 is the first mouse papillomavirus suitable for large-scale laboratory studies and has been proven to be highly malleable in our hands. We have shown that MmuPV1 has both cutaneous and mucosal tropisms (Cladel et al., 2017a; Cladel et al., 2017b; Cladel et al., 2013). For this study, we carried out two experiments with these mice. 1) Naïve animals were infected with infectious virions via the tail vein. As in the case of the rabbit model, infections developed at susceptible sites (both cutaneous and mucosal sites) in these intravenously infected mice. *Furthermore, active infections were detected in the stomachs of three of the tested animals.* 2) Blood was drawn from infected mice about seven month post infection and transfused into naïve animals. Once again, infections developed at medically relevant cutaneous and mucosal sites in the naïve animals. These sites included the mucosa of the vagina, penis, oral cavity and anus, as well as cutaneous tail and muzzle sites. Furthermore, active *infections were detected in the stomach of one of the tested animals*. These results are provocative and provide direct evidence of blood transmission of papillomavirus from an infected animal to a naïve animal. Our findings consequently call into question the safety of human blood supplies. They raise the question “Should donor blood be routinely screened for the presence of HPVs before being distributed to patients?”

## RESULTS

### Intravenous (IV) delivery of CRPV virions replicates the patterns observed with skin infections

If papillomavirus infections are capable of being transmitted via the blood, that implies the infectious agents are able to survive in the blood. We first applied the rabbit model to test whether CRPV virus deliberately introduced into the blood stream could yield infections at prewounded skin sites. Two outbred NZW rabbits (NZW#1 and NZW#2) and one HLA-A2.1 transgenic rabbit were used in this pilot study. Each animal was infected via the marginal ear vein with CRPV virions (500 µl of the viral stock= 2.75×10^10^ viral DNA equivalents) (Supplementary Table 1). The amount of virus in the circulation was estimated to be 1.8 ×10^5^ copies/µl blood. This figure is based on an estimated blood volume of 150 ml per animal. On each animal, eight back skin sites had been wounded three days prior to introduction of virus as per our local skin infection protocol (Cladel et al., 2008). The pre-wounded sites were scratched with a 28-gauge needle (Cladel et al., 2008) following intravenous (IV) viral infection and then monitored for tumor growth. Four weeks post infection, tumors were visible at the wounded sites of all three animals, and the tumors increased in size over time during the first six weeks post infection (Fig. 1 A-C). The tumor growth pattern mimicked the patterns observed in local skin infections in our previous studies (Hu et al., 2007b). Tumors in the transgenic animal (Fig. 1 C) regressed over time, as has often been observed in these animals with local skin infections (Hu et al., 2007b). Tumors on NZW#1 persisted as shown in Fig. 1 D. Interestingly, no tumors developed at the marginal ear vein sites, which further confirms previous findings that prewounding is crucial for viral infection in our CRPV/rabbit model (Cladel et al., 2008).

**Figure 1.**
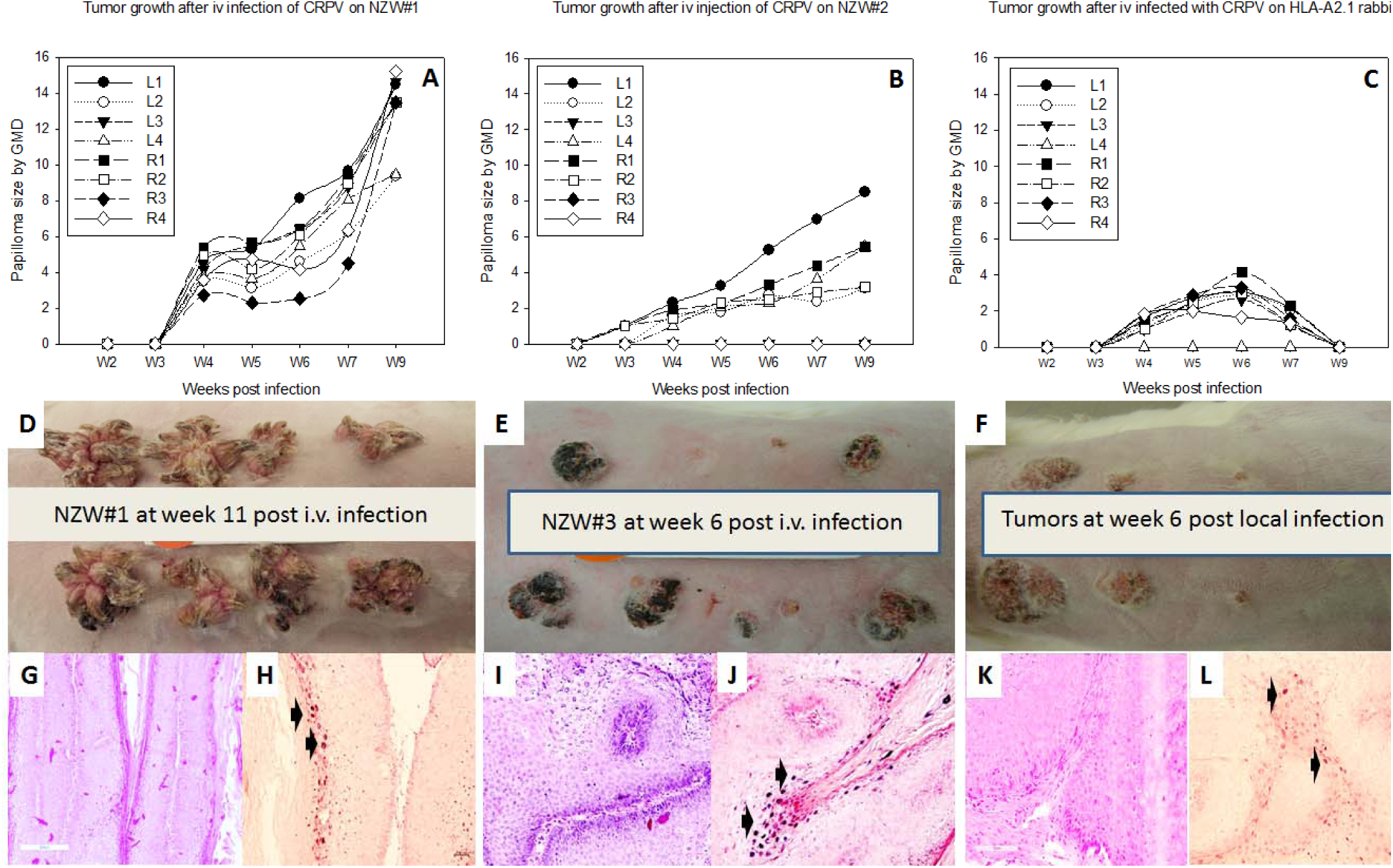
Tumor growth patterns resulting from IV infection with *CRPV virions* via marginal ear vein injection. **(A-C)** Two NZW rabbits (NZW#1-2) and one HLA-A2.1 transgenic rabbit (A2) infected by IV injection of virions equivalent to 2.75×10^10^ copies of viral DNA displayed tumors that mimic those by local skin infections in our previous studies (Hu et al., 2007b). **(D)** Tumor growth on one (NZW#1) of the three animals at week 11 post infection was shown. **(E)** one (NZW#3) of four additional rabbits (NZW#3-6) infected by IV injection of virions equivalent to 5.5×10^9^ viral DNA equivalents developed tumors at six weeks post infection. Both tumors (D and E) exhibited similar appearance of tumors from local skin infected with high to low dilutions of virus at week six post infection (**F**). **(G, I)** The tumors induced via marginal ear vein (IV) infection have similar morphology and histology (H&E, 20×) to those **(K)** initiated by local infections (H&E, 20×). (**H, J and L**) On the surface of the mass there was a subcorneal cleft filled with erythrocytes and fibrin with frequent heterophils (hemorrhagic vesicle), with scattered smaller acute stromal and intra-epithelial hemorrhages. No interface inflammation was detected in these tumors. Viral DNA was detected by *in situ* hybridization (ISH in tumors induced by both intravenous and local skin infections (20×, arrows).

The experiment was repeated using five-fold fewer virions (3.6 ×10^4^ /µl blood) to infect four naïve NZW rabbits (NZW#3-6) intravenously. In addition to eight prewounded sites, two back skin sites on each animal were wounded only on the day of IV infection. 18 out of 32 pre-wounded sites developed papillomas, whereas none of the eight sites wounded on the day of IV infection developed papillomas (P<0.05, Fisher’s exact test). These results recapitulate our findings for local skin infections (Fig. 1 F) and lend further support for the role of wound healing in the development of papillomavirus infections (Cladel et al., 2008). Again, no lesion was found at any IV injection sites on these rabbits.

We further compared the tumors induced by IV infection and those induced by local skin infection by histological analysis. Representative tumors from each of two IV infection experiments (Fig. 1 G and I, 20×) are shown. Histology similar to that seen in local skin infection (Fig. 1 K, 20×) is observed by H&E staining. Viral presence was detected by in situ hybridization (Fig. 1 H and J, 20× arrows) and is similar to that found in the rabbits with local skin infection (Fig. 1 L, 20× arrows).

Anti-CRPV antibodies were detected in the sera of all intravenously infected animals, but antibody level did not correlate with tumor size (Fig. 2 A). These antibodies were also neutralizing in our in vitro neutralization assays (Fig. 2 B). Collectively, these findings prove that CRPV circulating in the blood of rabbits can initiate infection at prewounded cutaneous sites, the preferred sites for CRPV infection, and stimulate anti-viral immune responses in these rabbits just as found in local skin infected animals (Hu et al., 2007a).

**Figure 2.**
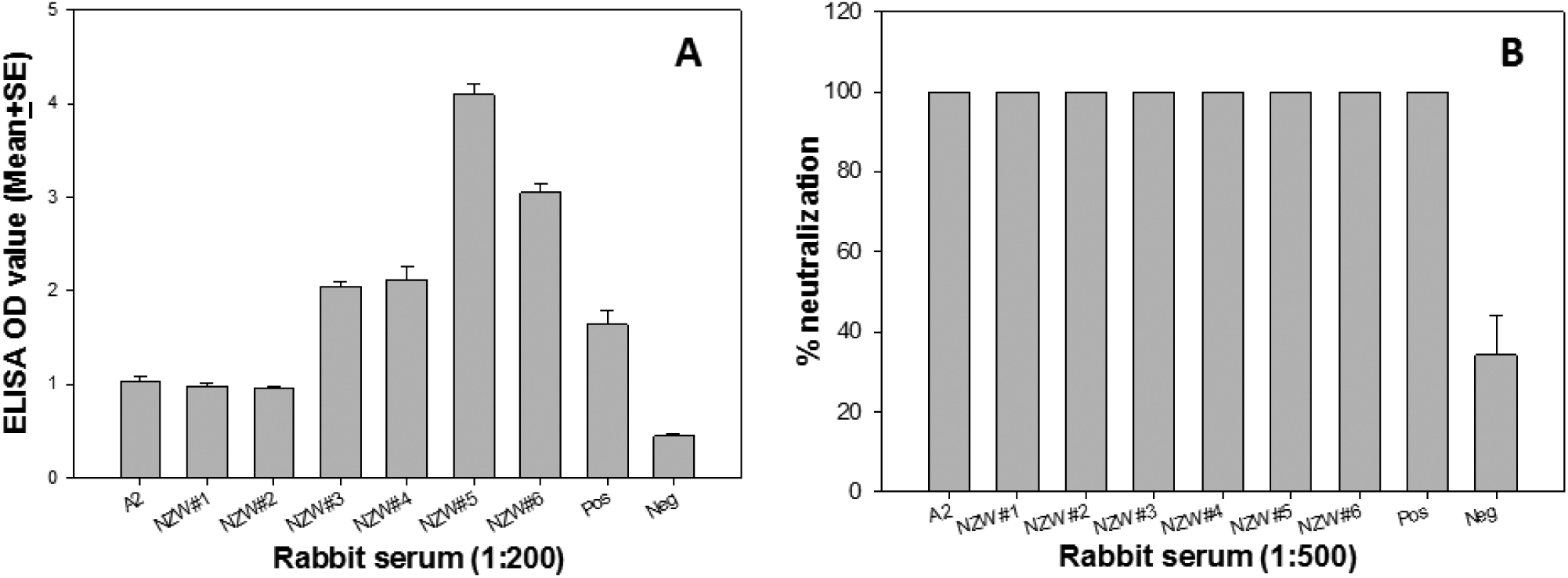
Neutralizing antibodies were induced in the intravenously infected rabbits. Positive control was anti-rabbit L1 (CRPV.1A) (Christensen and Kreider, 1991) and the negative control was serum of a non-infected rabbit for both ELISA and in vitro neutralization analyses. All seven IV infected animals generated specific anti-CRPV antibodies. **(A)** The antibody titer in ELISA assay was not correlated with the tumor size. **(B)** These anti-CRPV antibodies were neutralizing by the in vitro neutralizing assay.

### RNA sequencing (RNA-seq) analysis demonstrated similar CRPV transcription patterns for all cutaneous tumors resulting from either local skin infections or intravenous CRPV infections

To analyze CRPV transcripts arising from the tumors generated by CRPV virions through marginal ear vein injection, total RNA was isolated from four tumors from four different animals, depleted of ribosomal RNA and analyzed by RNA-seq. By mapping the RNA-seq raw reads to the newly arranged linear CRPV genome starting from nt 7421 and ending at nt 7420 using RNA sequence aligner TopHat, we obtained 18318, 24014, 62100, and 128869 viral reads for the respective tumor tissues. These reads account for 0.0290%, 0.0442%, 0.0911%, and 0.1960% of total RNA reads obtained from these samples. By uploading these uniquely mapped viral RNA reads to the Integrative Genomics Viewer (IGV) program to visualize reads coverage profile in parallel with the CRPV genome, we found three major coverage peaks in the E6, E7 and E1^E4 regions among all tumor tissues (Fig. 3).

**Figure 3.**
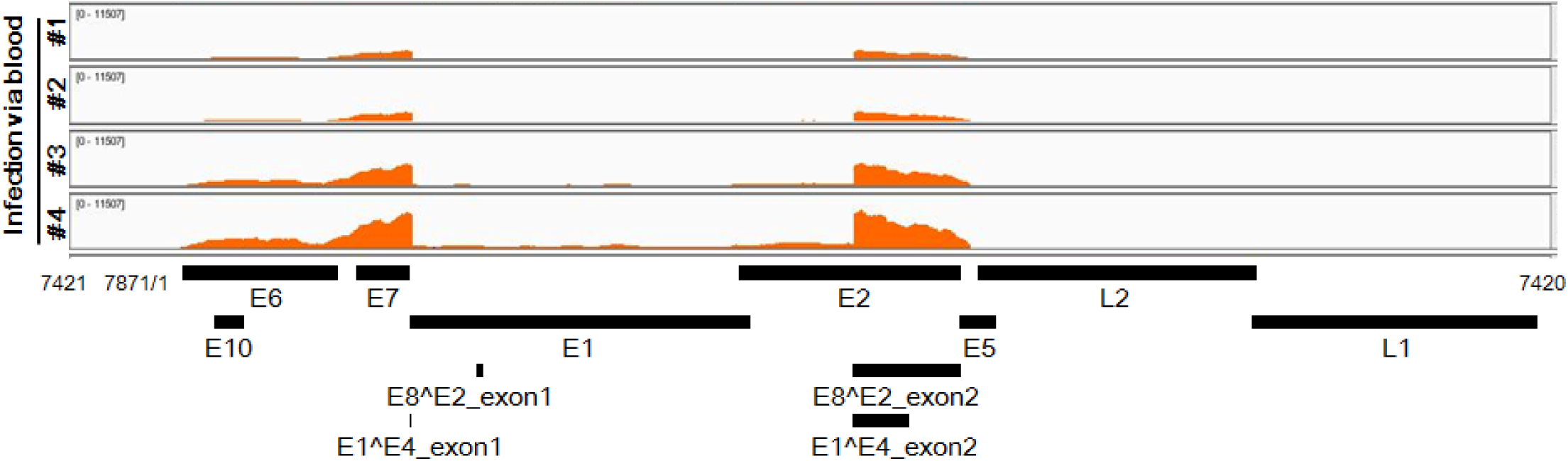
Viral RNA transcripts in four tumor lesions of four individual rabbits IV infected by CRPV virions through marginal ear vein injection. Total RNA isolated from each tumor and depleted of ribosomal RNA was analyzed by RNA-seq. By mapping the RNA-seq raw reads to the newly arranged linear CRPV genome starting from nt 7421 and ending at nt 7420 using RNA sequence aligner TopHat, we obtained 18318, 24014, 62100, and 128869 viral reads for the respective tumor tissues; this accounts for 0.0290%, 0.0442%, 0.0911%, and 0.1960% of total RNA reads obtained from each sample, respectively. By uploading these uniquely mapped viral RNA reads to the Integrative Genomics Viewer (IGV) program to visualize reads coverage profile along with the CRPV genome, we found three major coverage peaks in the E6, E7 and E1^E4 regions among all tumor tissues.

RNA-seq data of three groups of ten examined tissues (four normal skin tissues and three tumor tissues each induced by IV infection or local skin infection) were further analyzed by Principle component analysis (PCA). A well grouped dataset was found (Fig. 4 A). Interestingly, one of three tumors derived from IV infections had high virus titer by RNA-seq raw reads (Fig. 3) and the altered gene expression pattern was close to that of the tumors induced by local skin infection (Fig. 4 A). Volcano plots of 17742 annotated genes in the rabbit genome exhibited significant differences in the host transcriptome of the normal control skin group compared with the tumor groups induced either by IV infection or local skin infection (Fig. 4 B-C). The dysregulated expression of the genes is detailed in supplemental Table 3. Approximately 3,000 out of 5,224 genes were similarly dysregulated by the two routes of CRPV infections (Fig. 4 D). Overall, fewer genes (3,485) displayed the altered expression in IV infected tumors when compared with those in locally skin-infected tumors (4,799) (Fig. 4 D). Using the thresholds of P≤0.05 and absolute fold change (FC) ≥2.0 of differentiated gene expression, we analyzed the top 100 up-regulated and top 100 down-regulated genes in each experimental group (Fig. 4 F) by heatmap analysis and identified the most important genes with differential expression common in the tumors derived from both routes of virus infections (Fig. 4 E), notably the genes with upregulated expression. For the genes with reduced expression, five expression patterns could be subgrouped according to their reduced expression levels, with the expression of only a few genes being commonly downregulated in the tumors derived from both infection routes. A majority of them exhibited the expression reduction from cutaneous infection low to IV infection high or vice versa (Fig. 4 E). Based on gene function and its RNA abundance, we subsequently verified nine rabbit genes with significantly different expression in CRPV-induced tumors from both infection routes by real-time qPCR (Table 1) (Fig. 5 A-B). Consistent with RNA-seq data, the expression of SLN, TAC1, MYH8, and PGAM2 were down-regulated, whereas SDRC7, KRT16, S100A9, IL36G, and FABP9 were all up-regulated in both blood (Animal #9, #11, and #12) and skin (Animal #6, #7, and #8)-induced tumors when compared to those in normal skin of control animals (Fig. 5 B). Consistent with the RNA-seq results, western blot analysis of two selected genes further confirmed the increased protein expression of S100A9 and decreased expression of APOBEC2 in tumors induced by both routes of infections (Fig. 5 C). Taken together, these findings strongly support the hypothesis that bloodborne and local skin papillomavirus infections have similar infectivity mechanisms.

**Table 1.**
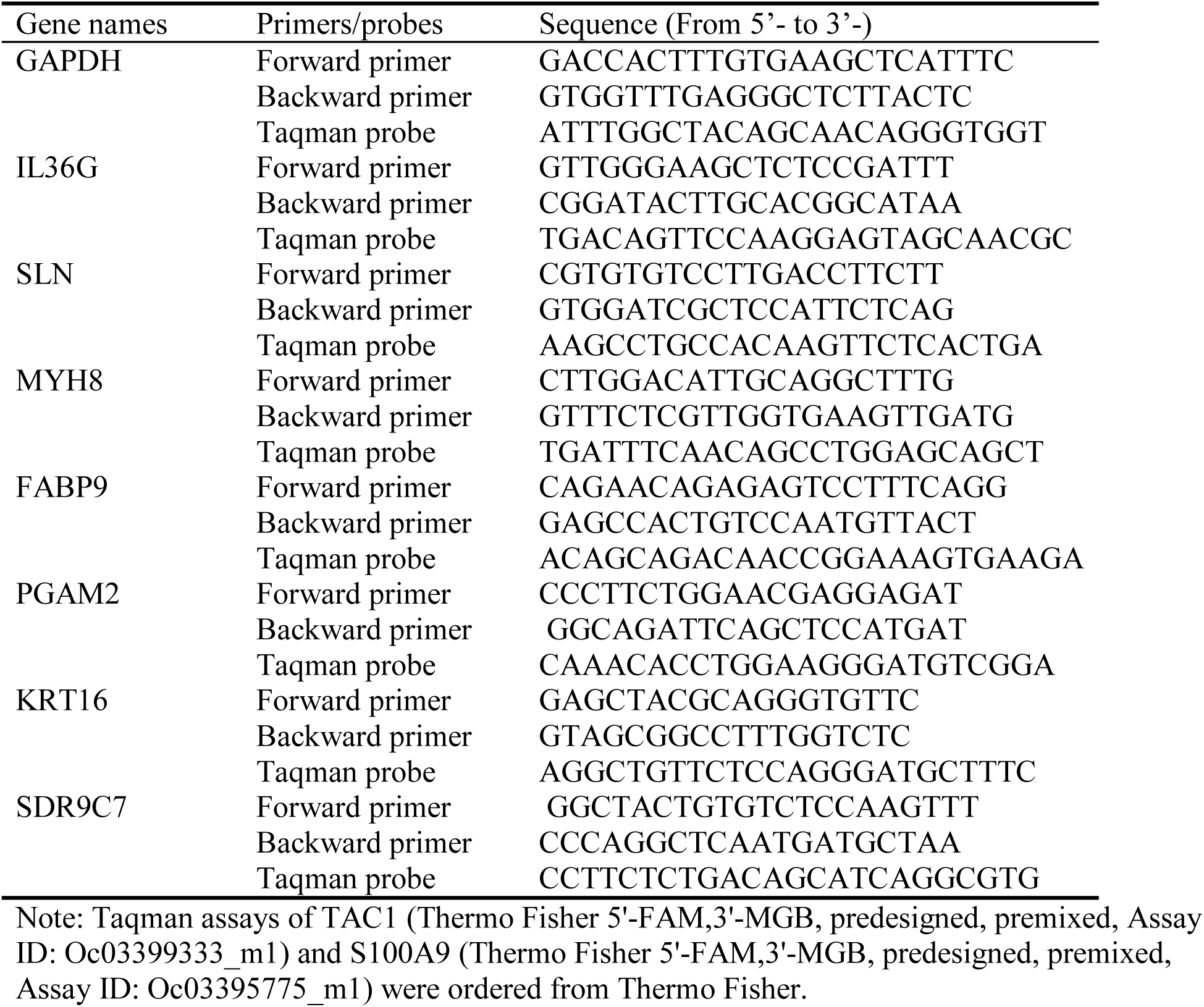
Primers and probes for RT-qPCR of rabbit gene transcripts.

**Figure 4.**
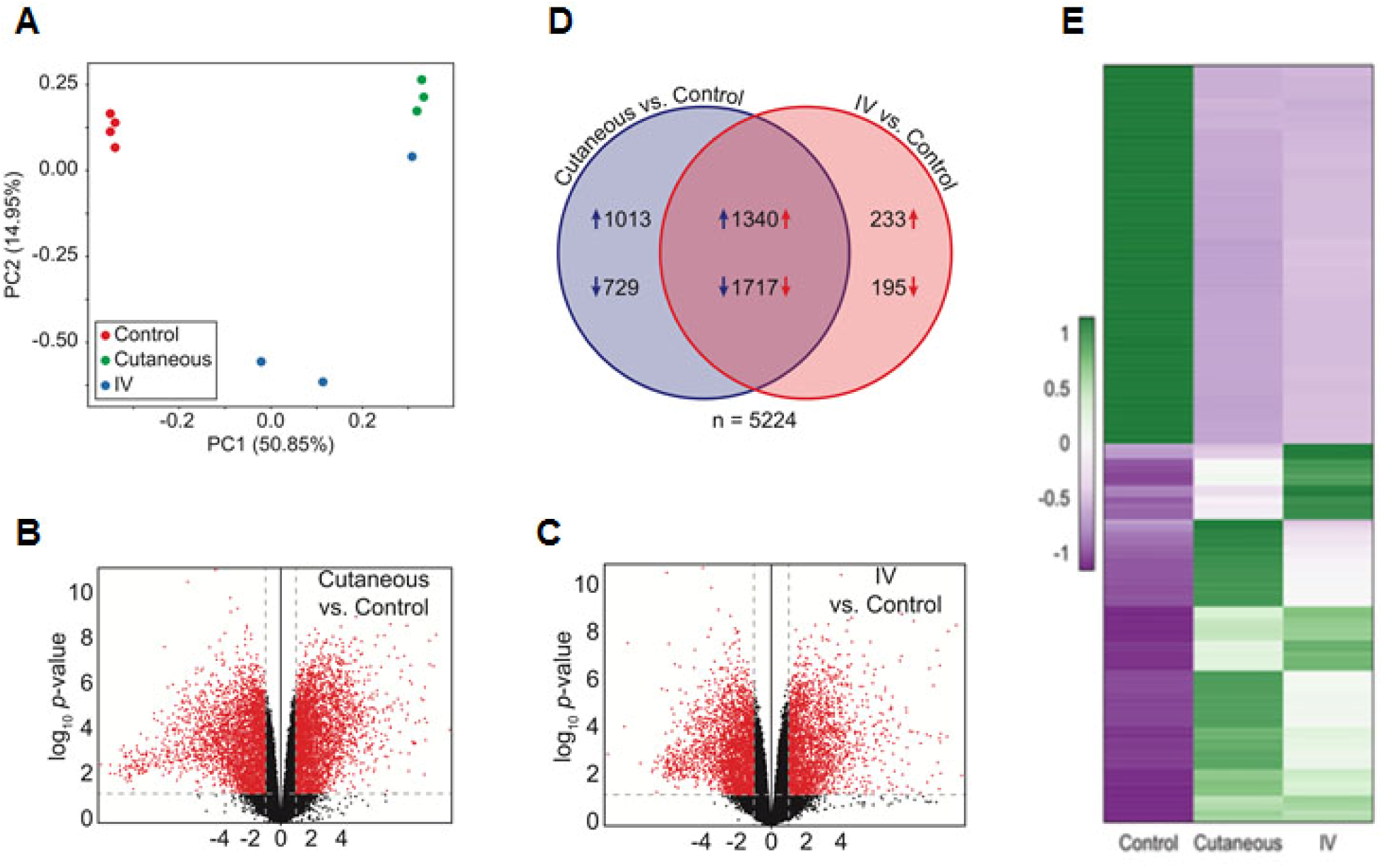
Dysregulation of the host transcriptome by CRPV infection. **(A)** Principle component analysis of the ten RNA-seq samples. **(B-C)** Volcano plots of 17742 annotated genes assayed in each contrast of Consistent with RNA-seq our analysis. The x-axis is the log2 fold change in expression (note the x-axes of each panel are not to the same scale). The y-axis is *p*-value adjusted for multiple comparisons. Red dots indicate the genes with both significant differential expression and large absolute fold change relative to control; black dots indicate those genes that do not met these criteria. Vertical dashed lines represent fold change thresholds (absolute fold change ≥ 2.0) and horizontal dashed lines represent the significance threshold (adjusted *p* ≤ 0.05). **(D)** Venn diagram of all 5224 genes with differential expression (adjusted *p* ≤ 0.05 and absolute fold change ≥ 2.0) in the wart tissues induced by local skin or IV CRPV infection over the normal control skin tissues. Numbers with arrows indicate the number of genes (after all filters) up- and down-regulated in each experimental group relative to control group. **(E)** Heat map showing the top 100 up-regulated and top 100 down-regulated genes with significantly differential expression common both in the tumors induced by both local skin and IV infections, relative to control. A color scale bar represents relative gene expression level within centered rows. Unit variance scaling has been applied to rows. Both rows and columns are clustered using Euclidean distance and complete linkage.

**Figure 5.**
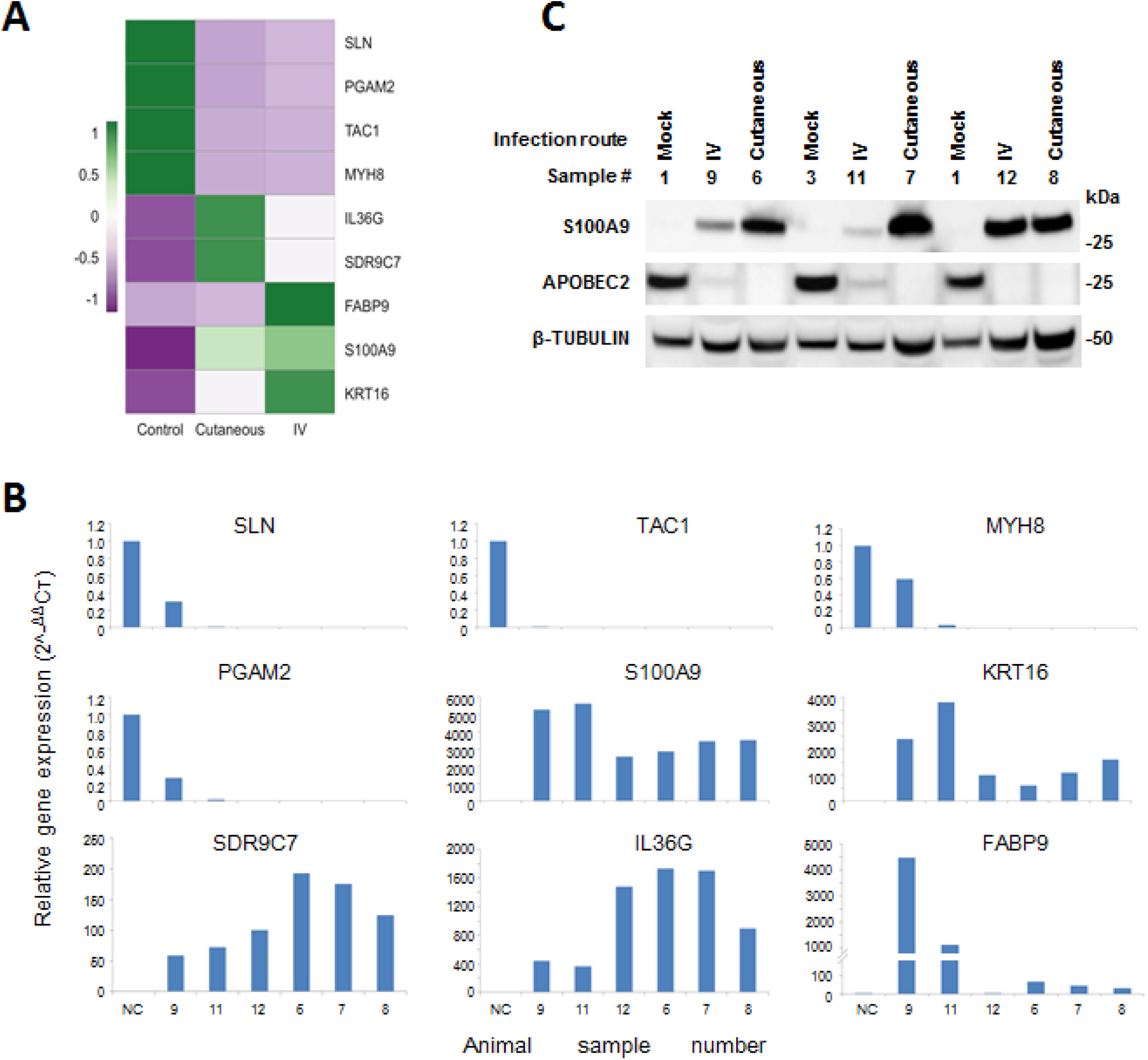
Representative host genes with differential expression in skin tumors induced by both routes of CRPV infections. **(A)** Heat map showing the expression of 9 selected host genes chosen based on their expression abundance and cellular functions. A color scale bar represents relative gene expression level within centered rows. Unit variance scaling has been applied to rows. Both rows and columns are clustered using Euclidean distance and complete linkage. **(B)** Verification of differentially expressed rabbit genes in RNA-seq results by real-time qPCR. Consistent with RNA-seq data, SLN, TAC1, MYH8, and PGAM2 were down-regulated in both CRPV blood infection (Animal #9, #11, and #12) and CRPV skin infection (Animal#6, #7, and #8) animals relative to those in normal control animals, whereas SDRC7, KRT16, S100A9, IL36G, and FABP9 were up-regulated in both CRPV skin infection and CRPV blood infection animals compared to normal controls. The Y-axis indicates relative gene expression levels calculated by 2-^ΔΔ^CT and the X-axis indicate the different samples. NC, gene expression in the normal tissue, was set to 1 after normalization to GAPDH. **(C)** Western blot analysis of representative samples from normal skin and warts induced by cutaneous or intravenous (IV) CRPV infection for the expression of S100A9 and Apobec2. Cellular β-tubulin served as loading control.

### CRPV DNA delivered intravenously induced tumors at prewounded sites and replicated the pattern previously observed with DNA locally delivered to pre-wounded sites

CRPV DNA delivered locally to wounded sites results in tumor growth (Cladel et al., 2008; Hu et al., 2002; Kreider et al., 1995). HPV DNA can be detected in human blood (Bodaghi et al., 2005a; Chen et al., 2009; Cocuzza et al., 2017; Jeannot et al., 2016). We hypothesized that the rabbit model could be used to determine if viral DNA in the blood could pose a potential risk for infection. Our previous work determined that about 1.3×10^10^ viral DNA equivalents would be needed to guarantee tumor growth in all skin sites locally treated with viral DNA, and that a dose as low as 1.3×10^9^ viral DNA equivalents would be capable of inducing tumors at a subset of sites (Supplementary Table 2). To accommodate for dilution as well as possible degradation of DNA in the circulatory system, we inoculated 500 µg of viral DNA (estimated to be 4.6×10^11^ copies /µl blood) into the ear veins of two rabbits with pre-wounded skin sites on the backs. Both rabbits grew tumors at these pre-wounded skin sites at week nine post IV infection. Histology (Fig. 6 B) of one representative papilloma (Fig. 6 A) was similar to that of the tumors initiated by local skin infections by virions (Fig. 1 K) with typical epithelial koilocytes in the infected area.

**Figure 6.**
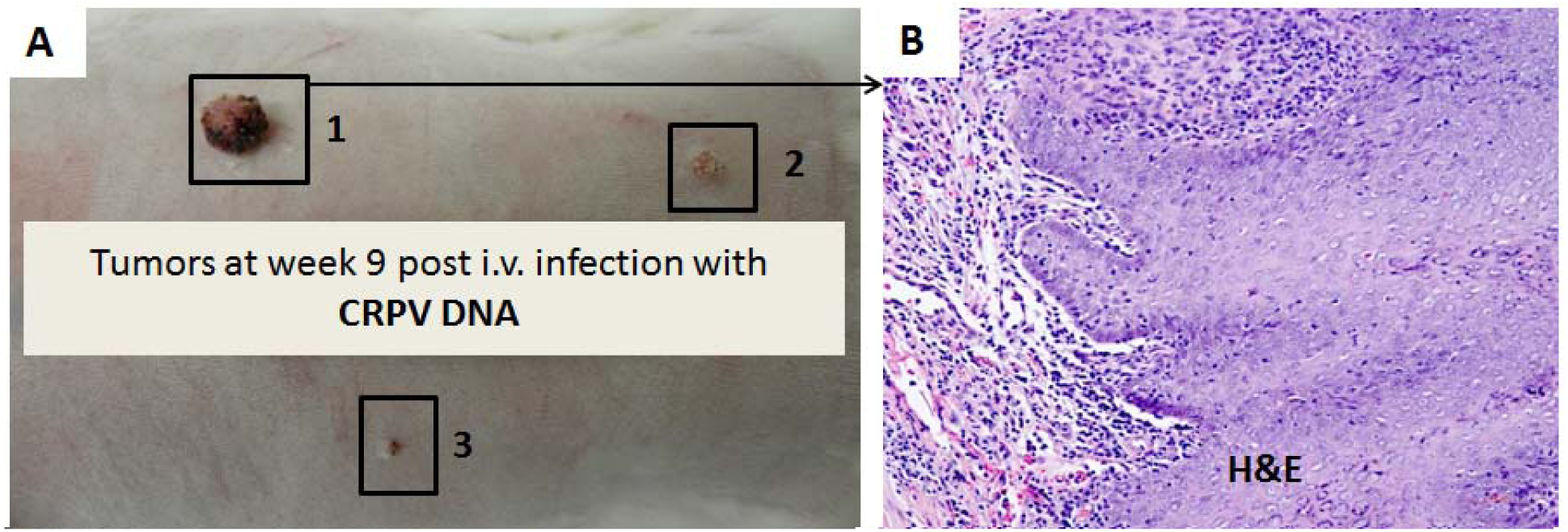
Tumor growth by IV infection with *CRPV DNA* via marginal ear vein. **(A)** Three tumors were detected at pre-wounded back skin sites on one rabbit at week nine post intravenously infected with CRPV DNA via the marginal ear vein injection. **(B)** There are multifocal dense leukocytic infiltrates at the dermal-epidermal junctions within the tumor mass, predominantly macrophages with fewer lymphocytes and rare neutrophils in a representative tumor (H&E, 20×). Epithelial polarization and differentiation are maintained from basement membrane to the surface, with occasional mitoses and mild nuclear atypia. There are occasional koilocytosis and scattered individual apoptotic keratinocytes within the epithelium. There is no invasion present in examined tissues. This tumor has similar histology to those initiated by local skin infection as shown in Fig. 1 K.

### Transfusion of blood containing CRPV virus resulted in tumors at prewounded back sites

The driving question behind our research was “Can transfusion of human blood containing HPV sequences result in papillomavirus infection in the recipient?” This question cannot be answered with experiments using human subjects, but it can be approached using animal models. In a small pilot study designed to provide proof of principle data, two outbred NZW domestic rabbits were intravenously infected with CRPV virions (1.8 ×10^5^ copies/µl blood). Twenty-five minutes after the virus inoculation, 10 mL of blood was withdrawn from each donor rabbit and infused into corresponding siblings, each with eight pre-wounded back skin sites. Viral concentration was estimated to be 1.3 ×10^4^ copies/µl of blood. One recipient grew a skin tumor ten weeks after receiving blood from the donor (Fig. 7 A). Histology of this tumor (Fig. 7 B) is similar to that shown in Fig. 1K. Viral DNA was detected in the tumor by in situ hybridization (Fig. 7 C, 60×). No tumors were detected at the sites that had not been pre-wounded.

**Figure 7.**
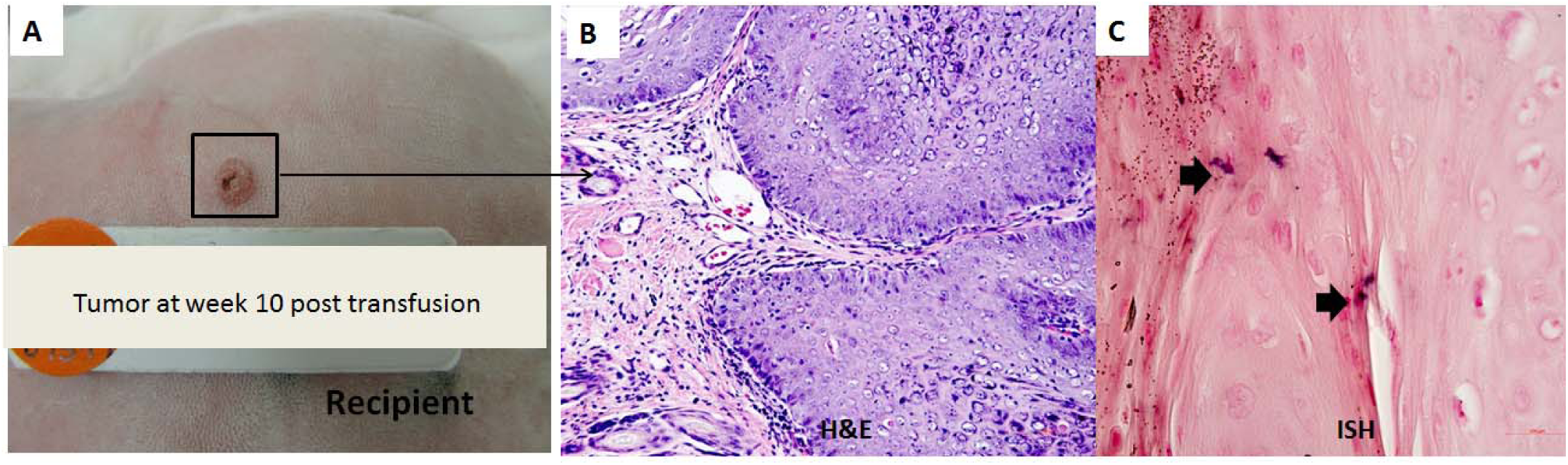
Tumor growth on blood transfusion recipient. **(A)** A tumor was detected at one recipient rabbit’s back pre-wounded skin site at week ten after receiving a blood transfusion from a rabbit that had been infected via the ear vein with 1.8 ×10^5^ viral DNA equivalents /µl blood for 30 minutes. **(B)** The tumor has exophytic polypoid cutaneous warty masses similar to Fig. 4B, but with minimal inflammation. There is a mixture of ortho- and parakeratotic debris on the surface of the masses. **(C)** Viral DNA was detected in the tumor by in situ hybridization (ISH, 60×).

### Athymic mice inoculated intravenously with MmuPV1 developed infections at both cutaneous and mucosal tissues

Following the demonstration that CRPV delivered intravenously could yield infections and tumor formation at prewounded skin sites, we next wished to determine whether the same would be true for athymic nude mice in which MmuPV1 infects both mucosal and cutaneous tissues (Hu et al., 2017). We were also interested in a possible sex bias during MmuPV1 infection. To explore this possibility, this experiment was conducted using equal numbers of male and female animals.

1×10^8^ viral DNA equivalent virions were carefully injected into the tail vein of six female (Fig. 8 A) and six male (Fig. 8 B) Hsd: NU mice that had been pre-wounded at both cutaneous and mucosal sites according to our standard protocol (Cladel et al., 2013; Hu et al., 2015). The injection tail vein sites were treated topically with an excess dose of neutralizing monoclonal antibody (MPV.A4) immediately post injection to neutralize any virions remaining at the site (Supplementary Figure 1) (Cladel et al., 2017b). The animals were monitored for tumor growth at the prewounded cutaneous sites and for viral DNA by qPCR at mucosal sites (Hu et al., 2015). All tails of the infected mice grew tumors at the prewounded sites (representative mouse from female and male groups respectively, Fig. 8 A, B). No tumors developed at the sites of injection during the first ten weeks, indicating MPV.A4 efficiently blocked any possible skin contamination during tail IV injection. Mucosal sites (the oral cavity, anus, vagina, and penis) were positive for viral DNA by qPCR (Hu et al., 2015), as well as by immunohistochemistry and in situ hybridization (ISH) (Fig. 8 C).

**Figure 8.**
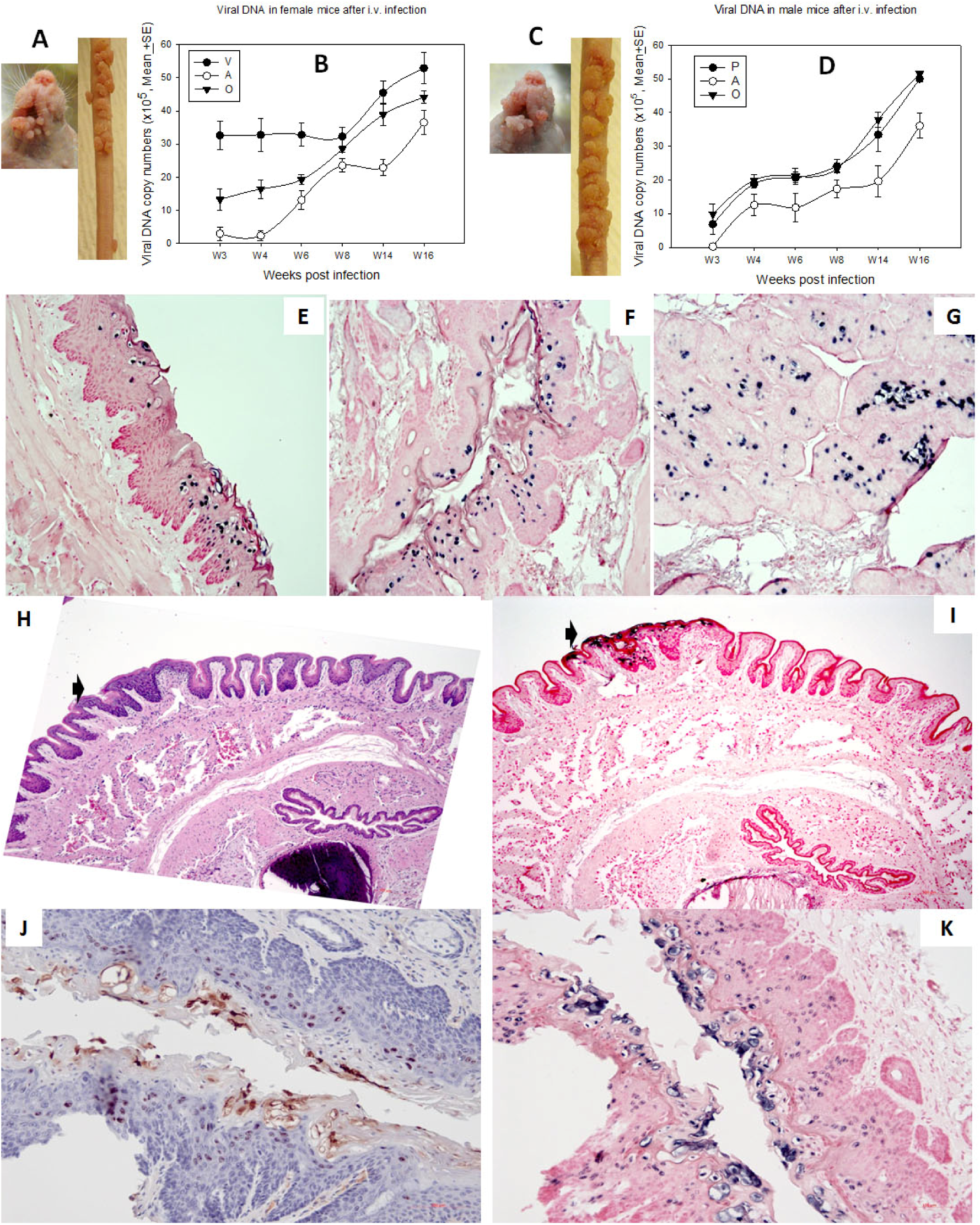
Tumor growth at cutaneous sites (muzzle and tail) and viral DNA detection at the four mucosal (vaginal, V; Penile, P; Anal, A and Oral, O) sites after MmuPV1 IV infection via the tail vein. Infections were introduced via the tail vein with 1×10^8^ viral DNA equivalent virions in eight Hsd: Nu female and six male mice that had been pre-wounded according to our standard protocol. The injection sites were treated topically with an excess of neutralizing antibody (MPV.A4) immediately post injection to neutralize any virions remaining at the site (Cladel et al., 2017b). (**A, C)** The animals were monitored for tumor growth at pre-wounded cutaneous sites and **(B, D)** for viral DNA detection by Q-PCR at mucosal sites. **(E)** Tongue, **(F)** Vaginal, **(G)** Anal, (all 20×), were positive for viral DNA by in situ hybridization (ISH). Interestingly, dysplasia was found in the penile tissue (H&E, H, 10×, arrow) that was positive for viral DNA by ISH (**I**, 10×, **K**, 20×) and viral capsid protein by immunohistochemistry **(J,** 20×).

### Active infections were found in stomach tissues of mice infected via tail vein injection with MmuPV1

In view of the many observations in the literature that papillomavirus sequences can sometimes be found in cancers of internal organs, we examined whether blood-borne infections could result in internal organ infections. The organs of selected intravenously infected mice were examined for tissue pathological changes by H&E staining; for viral DNA by in situ hybridization; and for viral capsid protein L1 by immunohistochemistry. Interestingly, the non-granular stomach tissues of three out of eleven intravenously infected mice were found to be positive for both viral DNA and L1 protein (Fig. 9). One of the three stomach tissues showed a focally extensive plaque lesion of mild to moderate hyperplasia with atypia (Fig. 9 A). Abundant hyperkeratosis with what appeared to be crypt formation was also found. There were occasional positive nuclei within the stratum spinosum and granulosum with abundant positive staining of both viral DNA and viral L1 protein in the cornified layers. There were multifocal cytoplasmic hybridization signals with parietal cells in the glandular stomach. Two stomach tissues displayed a small isolated (possibly pedunculated) focus of mild hyperplasia with cytological and nuclear atypia (low grade) in the non-glandular stomach with scattered individual nuclear positive cells (Fig. 9 D). These tissues were positive for both viral DNA, shown by in situ hybridization (Fig. 9 B and E, 20×, in blue), and for viral capsid protein, shown by immunohistochemistry (Fig. 9 C and F, 20×, in red). No other organs were found to be positive for MmuPV1 in the examined animals. This observation indicates that mouse tissues not normally permissive for MmuPV1 infection can become infected when the route of viral delivery is via the blood.

**Figure 9.**
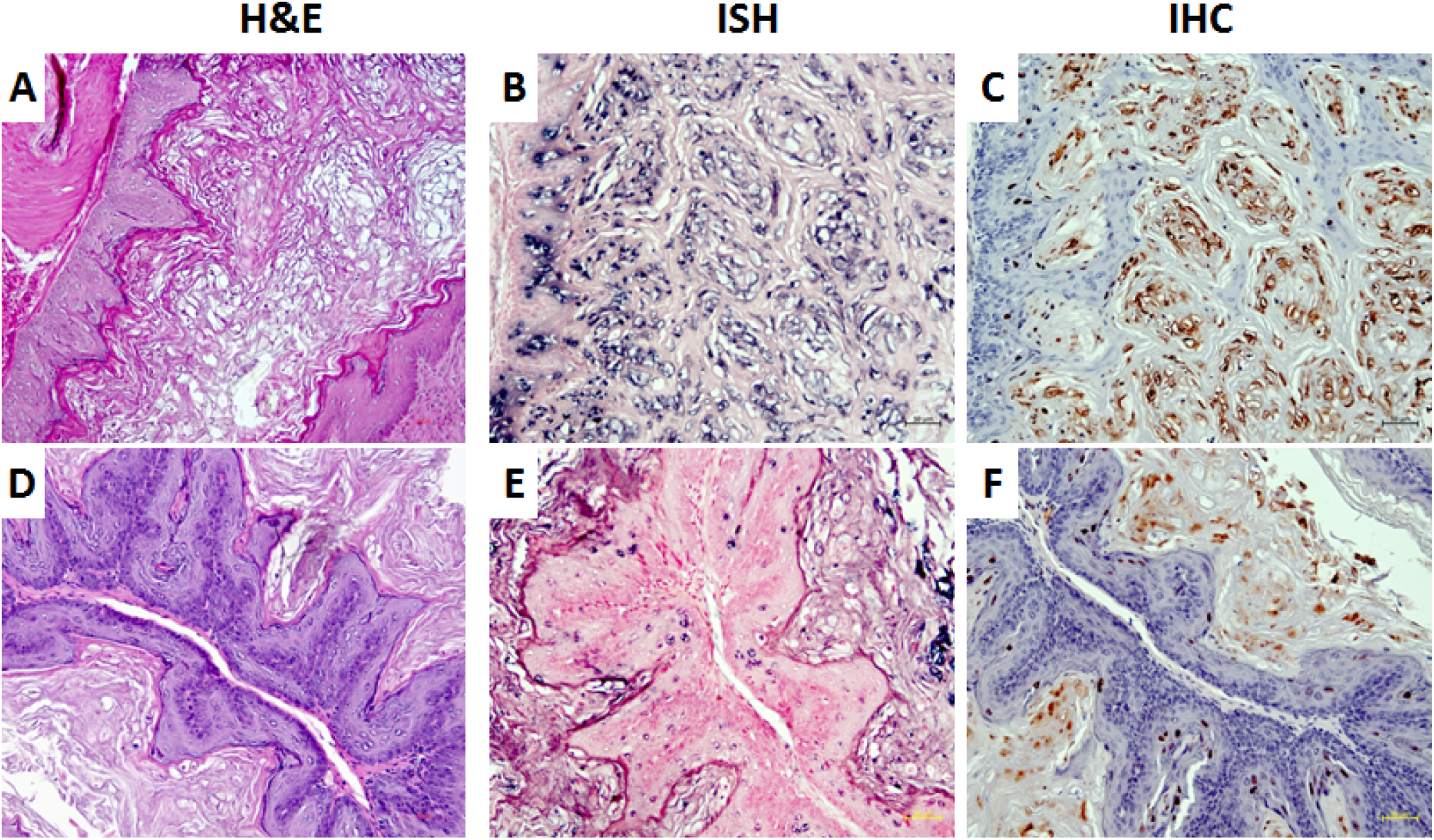
Three (two females and one male) of eleven tested mice were positive for virus infection in the stomach tissues. Representative stomach tissues were examined for histology. **(A)** Within the glandular stomach there is a focally extensive plaque lesion of mild to moderate hyperplasia with atypia. There is abundant hyperkeratosis (likely parakeratotic), with what appear to be crypt formation. There are occasional positive nuclei within the stratum spinosum and granulosum, with abundant positive staining of the cornified layers. There is multifocal cytoplasmic hybridization with parietal cells in the glandular stomach. **(D)** A small isolated (possibly pedunculated) focus of mild hyperplasia with cytologic and nuclear atypia (low grade) in the non-glandular stomach with scattered individual nuclear positive cells. These tissues were positive for viral DNA by both *in situ* hybridization (ISH, **B, E,** 20×, in blue) and viral capsid protein by immunohistochemistry (IHC, **C, F**, 20×, in red).

### Transfusion of naïve mice with blood from mice with active infections yielded tumors at prewounded sites

Since the impetus for this study was the concern that papillomaviruses might be spread from an infected to an uninfected individual via the blood, we next examined the ability of mice with active infections to transmit the infection to naive animals via blood transfusion. Two infected mice (one male and one female) were sacrificed at seven months after initial IV infection and 0.2 ml of blood from each animal was transfused via tail vein injection into three male (M1-M3) and three female (F1-F3) naïve littermates. The injection tail vein sites were treated topically with an excess dose of neutralizing monoclonal antibody (MPV.A4) immediately post injection to neutralize any virions remaining at the site (Cladel et al., 2017b). The corresponding recipients had been pre-wounded according to our usual protocol (Cladel et al., 2017a). At ten weeks post infection, all recipients, both male and female, were positive for viral infection and tumor growth at all of the wounded sites (Fig. 10 A-D). No lesions developed at the sites of injection. The tissues were positive for viral DNA (Fig. 10E). *Importantly, one stomach tissue was found to be positive for viral DNA* (ISH, 60×, in blue) and capsid protein (IHC, arrows, 60×, in red, Fig. 10 F). These findings conclusively demonstrate that blood from animals with papillomavirus infections can in fact transmit infections to naïve animals, especially those that are immunosuppressed. Furthermore, the stomach can become infected under these conditions.

**Figure 10.**
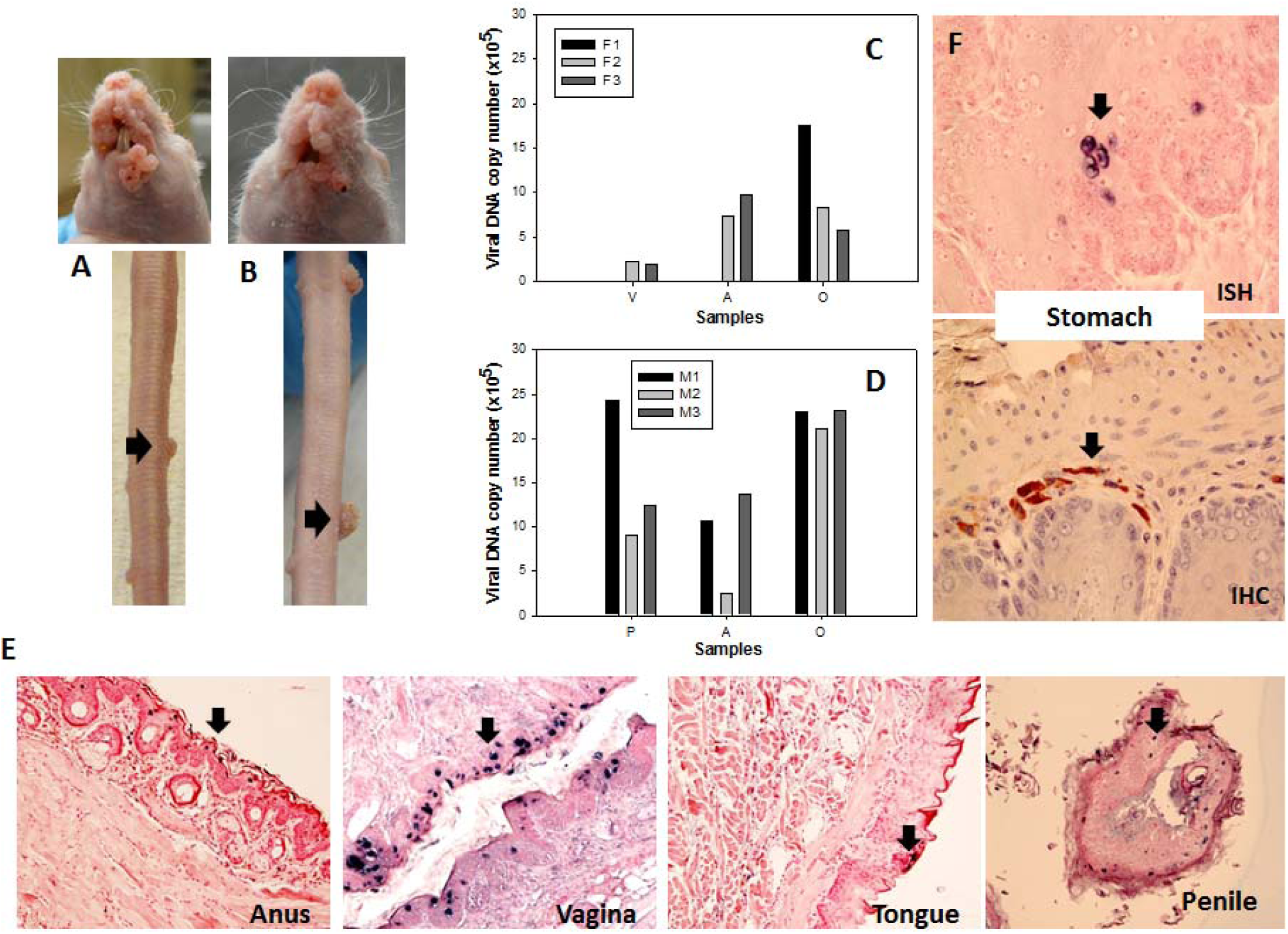
Blood of MmuPV1 infected mice with skin tumors was infectious at seven months post initial IV injection. Each naïve littermate transfused by IV injection of 0.2 mL of blood from two infected mice sacrificed seven months after initial IV MmuPV1 infection was examined weekly for tumor growth at the pre-wounded skin area. (**A, B**) Representative tumor growth (arrows) at the muzzle and the tail of naïve Hsd: Nu female (**A**) and male (**B**) mice at week sixteen post blood transfusion. Viral DNA was detected at the vaginal (V), Anal (A) and Oral (O) sites in three females (**C**, F1-F3) and the Penile (P), Anal (A) and Oral (O) sites in three male (**D**, M1-M3) mice by qPCR. Mucosal sites of these mice (Vagina, Anus, Tongue, and penile) were positive for viral DNA by in situ hybridization **(E**, arrows, 20×, in blue). One of the females was positive for viral DNA (ISH arrows, 60×, in blue) and viral capsid protein L1 (IHC, arrows, 60×, in red) in the stomach tissues **(F)**.

### Viral DNA was detected in blood samples of CRPV infected rabbits and MmuPV1 infected athymic mice

If blood transmission is one route of papillomavirus dissemination, then there should be a DNA signature in the blood of infected animals. PCR amplification, rolling circle amplification and DNA sequencing were performed to evaluate the presence of CRPV or MmuPV1 DNA in the blood of both locally and IV infected rabbits and athymic mice respectively. Viral DNA was detected in the whole blood of both CRPV-infected rabbits (12 /29) and MmuPV1-infected mice (8/12) which were used in our previous studies (Cladel et al., 2008). 12 out of 31 mouse serum samples also tested positive for the presence of viral sequences but none of the rabbit sera were positive. These findings suggest the possibility that systemic papillomavirus infections might have been established for much longer periods in the immunodeficient mice when compared with the immunocompetent rabbits. Consistent with these findings, a systemic infection has also been reported in bovine papillomavirus (BPV)-infected animals (Sadeghi et al., 2017).

## DISCUSSION

In this study we examine the significance of papillomavirus infection in the blood. If papillomaviruses can, indeed, be transmitted via the blood, millions of patients could be at risk of HPV infections (Laffort et al., 2004; Shanis et al., 2018). Blood is routinely screened for human immunodeficiency virus (HIV), hepatitis C (HCV), and hepatitis B (HBV). Procedures are being developed to detect emerging viruses such as Dengue and Zika (Stramer, 2014). However, there is no screening for HPVs. This omission appears to stem from the assumption that HPV-associated human cancers, primarily cervical cancer, are strictly sexually transmitted (Khoury et al., 2018). While that is true in many cases, it does not explain the finding that HPV can be detected in the blood of a subset of sexually naïve children with hemophilia who have received multiple transfusions (Bodaghi et al., 2005a). Nor does it explain the occasional presence of HPV sequences in tumors of organs not normally exposed to the virus via sex, such as the stomach, prostate, breast, colon, bladder, esophagus, and lung (Agalliu et al., 2018; Akhtar and Bansal, 2017; Damin et al., 2007; Malhone et al., 2018; Mirzaei et al., 2018; Russo et al., 2018; Salyakina and Tsinoremas, 2013; Shigehara et al., 2014). We asked whether such observations could be due to transmission of the virus through the blood.

All papillomaviruses are highly species specific. Therefore a human papillomavirus cannot infect any animal (Campo, 2002). As a result, studies of these viruses have historically been undertaken using animal models (Christensen et al., 2017). These include Cottontail Rabbit Papillomavirus (CRPV) and mouse papillomavirus (MmuPV1) (Ingle et al., 2011). Our laboratory has done much pioneering work with the CRPV/rabbit model and the MmuPV1/mouse model (Christensen et al., 2017; Hu et al., 2017). The domestic rabbit (*Oryctolagus cuniculus*) is not the natural host for CRPV. CRPV was isolated from its natural host, the wild cottontail rabbit of the Western United States (S*ylvilagus floridanus*) (Escudero Duch et al., 2015). While the tumors in both animals are similar in appearance, the cottontail rabbit produces far more virus than does the domestic rabbit (Christensen et al., 2017). Intriguingly, these tumors are highly localized and cross contamination has never been seen in our domestic rabbit CRPV studies. The MmuPV1 mouse model has been recently established and immunocompromised mice are most susceptible to MmuPV1. MmuPV1 secondary infections are common although secondary infections always appear much later than the primary infections (Cladel et al., 2017b; Cladel et al., 2013; Hu et al., 2015). By using a neutralizing monoclonal antibody MPV.A4 that completely neutralized both cutaneous and mucosal MmuPV1 infections when the athymic mice were passively immunized (supplementary Figure 1), we demonstrated that an excess of this antibody, immediately applied to the tail vein injection site after infusion with MmuPV1, could prevent contamination from IV infection. No tumors at these injection sites were ever observed.

Armed with the CRPV and MmuPV1 preclinical models, we determined to address the question of transmissibility of papillomaviruses through the blood. We demonstrated that 1) Viral sequences could be detected in the blood of infected animals; 2) Virus introduced into the blood could yield tumors at receptive sites, both cutaneous and mucosal; 3) CRPV DNA introduced into the blood stream yielded papillomas at prepared skin sites; 4) Similar mechanisms are used for infections via the blood and by direct application of virus to the skin as determined by RNA-seq analysis; 5) Transfusion of blood from an animal that had received virus via intravenous infection to a naïve sibling resulted in papillomas in the transfusion recipient; 6) Virus introduced via intravenous delivery not only resulted in infections at the expected sites but also at sites that are not normally permissive for MmuPV1 infections, and 7) Blood from animals with active infections could induce infections in naive mice when transfused into these animals.

We conducted RNA-seq analysis to compare the transcriptomes of CRPV tumors induced by local skin infection or IV infection. To allow for viral dilution in the blood during IV infection a 10,000 fold excess of virus was used relative to that used for local infection. Despite the possibility of stimulating host immune responses (Christensen et al., 1996), these IV infections induced tumor growth at the prewounded back skin sites. Interestingly, the total percentage of viral transcripts was much lower in tumors induced intravenously relative to tumors induced locally. This is most likely because only a small portion of the IV inoculated virus particles could reach the susceptible skin sites through the circulation. However, the patterns of viral transcription in the tumors were identical for both routes of infections. We further analyzed the host gene transcriptome of tumors resulting from both infection routes. Consistent with findings for the viral RNA reads, we found dysregulated expression of fewer genes in IV infection-induced tumors than in local infection-induced tumors. Although these two different infection routes induced different numbers of genes with altered expression, we found that the majority of the genes with significantly different expression were common to all tumors examined by RNA-seq. A subset of those with altered expression was selectively verified by RT-qPCR and Western blotting. These data provide further evidence that mechanisms for papillomavirus IV infection and local infection are similar.

Most individuals acquire papillomavirus infections at some point in their lifetimes (Gravitt and Winer, 2017). These infections are generally asymptomatic and the individual may not even know he or she is infected. Sexually transmitted infections are very common in young people, a cohort commonly represented in the pool of blood donors. Most papillomavirus infections are thought to clear spontaneously but the process often occurs over a lengthy time period of 1-2 years or more (Shew et al., 2015). In 10-20% of that HPV infected population, infection with the same viral species reappears at a later date (Martinez and Troconis, 2014). This supports the concept of latency as an alternative to clearance (Gravitt, 2011; Gravitt, 2012; Gravitt and Winer, 2017). In a small percentage of infected people, the viral infection does not clear at all and eventually progresses to cancer. Papillomaviruses are almost always implicated in cervical cancer, a leading cause of death in women of child bearing age in the developing world (Serrano et al., 2017).

Papillomaviruses are being identified in an increasing number of oral cancers, one of the malignancies that is rapidly escalating worldwide (Taberna et al., 2017). For reasons that are still unclear, the immune system is not capable of eradicating a subset of infections. It is possible that virus from acute or latent infections makes its way, in some yet to be determined manner, into the blood stream (Moustafa et al., 2017). Our results from animal studies would support this. Transfusion of this blood into another individual, especially one with a compromised immune system, could pose a risk of infection to the recipient. These infections might manifest not only in the genital sites normally associated with the virus but also in distant vital organs, as shown in this paper where the virus was found in the stomachs of four different animals. We postulate that blood transmission may be one mechanism whereby papillomaviruses reach these organs and subsequently contribute to cancer development.

To investigate the mechanism by which the virus and viral DNA are transported to local tissues and induce infections therein, we tested viral binding and infection in PBMCs harvested from rabbit blood. We observed the attachment of both virions and viral DNA to PBMCs, but detected no viral transcripts after three days of incubation, indicating no active infection occurred in these cells. These findings agree with reports in the literature that PBMCs are capable of binding and transporting virus via the circulatory system (Bodaghi et al., 2005a). Further studies will be needed to determine the specific blood monocytes preferred for viral attachment.

The link between a blood transfusion and a cancer that manifests much later in life would not be easy to detect in hindsight. In view of the results presented, it would seem prudent to screen blood routinely for HPV and to take steps to eliminate the pathogens when found. However, many questions remain unanswered. Among them: 1) Is there a threshold viral load beyond which blood-borne virus does not constitute a threat? 2) Are certain blood products free of virus? We have shown that PBMCs bind both virus and viral DNA. Other studies have demonstrated that viral DNA can be detected in serum and plasma (Cocuzza et al., 2017; Jeannot et al., 2016). We have detected viral DNA in serum of the infected mice. However, we have not yet investigated other blood products. 3) Is there an effective way to eliminate PV from donated blood? These and other questions will be the focus of future studies. It is not necessary to answer them, however, to conclude from the data generated to date that the presence of viral sequences in the blood could pose a long-term risk for patients receiving transfusions of whole blood or specific blood components. Therefore, we recommend the screening of all blood for the presence of papillomavirus sequences until such time that it can be proven that the presence of HPV does not pose a risk.

In summary, we have demonstrated, using two different animal models, that papillomaviruses can be transmitted through the blood and become infectious in recipients. Furthermore, transfusion of blood from an animal that had received virus via intravenous infection to a naïve sibling resulted in papillomas in the transfusion recipient. Finally, we have demonstrated in the mouse model that virus delivered via blood can produce active infections in the stomach. These results are provocative and call into question the serious implications of HPV in the human blood supply. Our findings suggest that the human blood supply could be a potential source of HPV infection. They further hint at a way that papillomaviruses might come to infect internal organs and set the stage for the development of cancers.

## MATERIALS AND MATHODS

### Animals and viral infections

All rabbit and mouse work was approved by the Institutional Animal Care and Use Committee of Pennsylvania State University’s College of Medicine (COM) and all procedures were performed in accordance with the required guidelines and regulations. Outbred New Zealand White (NZW) rabbits were purchased from Robinson and maintained in our animal facilities. HLA-A2.1 transgenic rabbits based on outbred NZW background were developed and maintained in our animal facilities (Hu et al., 2007c). The rabbits were housed individually. Male and female Hsd: NU outbred nude mice (Foxn1nu/nu) (6-8 weeks old) were obtained from ENVIGO. All mice were housed (2-3 mice/cage) in sterile cages within sterile filter hoods and were fed sterilized food and water in the COM BL2 animal core facility.

### Viral stock

Infectious virus was isolated from cottontail rabbit tumors or tumors on the tails of mice from our previous study (Cladel et al., 2016; Hu et al., 2007a). In brief, papillomas from the cottontail rabbits and tumors scraped from the tails of the mice were homogenized in phosphate-buffered saline (1×PBS) using a Polytron homogenizer (Brinkman PT10-35) at highest speed for three minutes while chilling in an ice bath. The homogenate was spun at 10,000 rpm and the supernatant was decanted into Eppendorf tubes for storage at −20ºC. For these experiments, the MmuPV1 was diluted 1:5 in 200 μl of 1×PBS and was passed through a 0.2 μm cellulose acetate sterile syringe filter. Viral DNA extracted from 5 μl of this virus stock was quantitated using qPCR as described (Hu et al., 2015).

### Titration of infectious CRPV virus and viral DNA

Viral skin infections can be initiated with either infectious virions isolated from the infected tissues of wild cottontail rabbits (Cladel et al., 2008) or viral DNA cloned into a plasmid vector (Kreider et al., 1995; Xiao and Brandsma, 1996). Our earlier work demonstrated the benefit obtained by wounding prior to infection and supported the theory that wounding is an important prerequisite for papillomavirus infection (Cladel et al., 2008). To identify the optimal concentration for both virions and viral DNA infections in the New Zealand White (NZW) domestic rabbits for the current study, titration studies were conducted using the pre-wounding skin infection method developed in our laboratory. Titration results demonstrated that more than 2.75×10^6^ virion DNA equivalents (Supplementary Table 1) and 1.3×10^10^ copies for cloned infectious CRPV DNA (Supplementary Table 2) are required to guarantee 100% tumor appearance at locally infected sites. However, as few as 2.75×10^3^ virion DNA equivalents or 1.3×10^9^ copies of the cloned infectious CRPV DNA were sufficient to generate clinical infections and papillomas at a subset of pre-wounded sites. No visible tumors could be detected if doses of virions or infectious viral DNA below these thresholds were delivered locally in prewounded sites. For the current study, 500 µl of infectious virus stock (2.75×10^10^ virion DNA equivalents) and 500 µg of the cloned infectious CRPV DNA (estimated to be 4.6×10^11^ copies /µl blood) were used for IV infection to maximize the chances for disease development in this model.

### Routes of infection and sample collection

CRPV infectious virions used in the current study were from a viral stock previously reported (Hu et al., 2014). CRPV DNA was cloned into a pUC19 vector as described in our previous publications (Hu et al., 2009). Animal back skin sites were pre-wounded three days before infection using our established pre-wounding techniques (Cladel et al., 2008a). For local infections, the rabbits were challenged with either virions or viral DNA at the pre-wounded sites (Hu et al., 2007b). To investigate whether intravenous (IV) infection with virions (500 µl of the viral stock= 2.75×10^10^ viral DNA equivalents) and viral DNA (500 µg of plasmid =4.6×10^11^ copies /µl blood) could induce tumors at distant susceptible back sites, six to eight back skin sites of different groups of rabbits were shaved and pre-wounded with a scalpel blade as previously described (Cladel et al., 2008) three days prior to IV infection. Virus or viral DNA was delivered via the marginal ear vein and then back skin sites were gently re-wounded with a 28G needle (Cladel et al., 2008). The infected animals were monitored for tumor growth weekly and tumors were measured and documented photographically. Rabbits were euthanized up to 12 weeks after initial viral infection, and tissues were collected for cellular, molecular, and histological analyses. Blood samples were collected from the intravenously infected rabbits at different time points. DNAs extracted from the whole blood using a blood DNA extraction kit from QIAGEN were used for viral DNA detection. Serum samples were harvested for the detection of anti-viral antibodies (Hu et al., 2007a). Peripheral blood mononuclear cells (PBMCs) were isolated from the blood samples and tested for viral sequences (Hu et al., 2010b). PBMCs were also harvested from naïve rabbits for in vitro binding and infection studies. To infect PBMCs, 5µl of CRPV virions or 1µg of viral DNA were incubated with 1×10^6^ rabbit PBMCs in vitro for 3-4 days at 37°C in RPMI1640 complemented with 10%FBS (Hyclone). The RNA was extracted and examined for the presence of viral transcripts as previously described (Hu et al., 2007a). To investigate whether transfusions of blood from an animal that had received virus (500 µl of the viral stock = 2.75×10^10^ viral DNA equivalents) by IV injection could transmit infection to naive siblings, 10 ml of blood was drawn from the injected animal heart 25 minutes post IV infection and transfused via IV injection into naive animals with pre-wounded back sites. Recipients were monitored for tumor growth and measurements were recorded. Rabbits were euthanized up to 12 weeks after viral infection, and tissues were collected for cellular, molecular, and histological analyses.

In addition to the CRPV/rabbit model, we used the more recently established MmuPV1/mouse model for a number of our studies (Hu et al., 2017). Female mice were subcutaneously inoculated with 3 mg Depo-Provera (Pfizer) in 100 µl PBS three days before the viral infection as previously described (Cladel et al., 2015). Mice were sedated intraperitoneally with 0.1ml/10 g body weight with ketamine/xylazine mixture (100 mg/10 mg in 10 ml ddH_2_O). The lower genital (vaginal) and anal tracts were wounded with Doctors’ Brush Picks coated with Conceptrol (Ortho Options, over the counter) as previously described (Hu et al., 2015). Tongues were withdrawn using a sterile forceps and microneedles were used to wound the ventral surface of the tongues (Cladel et al., 2016; Hu et al., 2015). Twenty-four hours after wounding, eight female and six male Hsd: Nu athymic mice were again anesthetized and challenged with infectious MmuPV1 virions (1×10^8^ viral DNA equivalents) via the tail vein. The injection sites were topically treated with neutralizing antibody (MPV.A4) immediately post injection to neutralize any virions remaining at the injection site. Monitoring was conducted weekly for infection at muzzle and tail and progress was documented photographically at these two cutaneous sites for each animal (Cladel et al., 2017a; Cladel et al., 2017b). Viral infection at three mucosal sites (vagina or penis, anus and tongue) was monitored for viral DNA using swabs or by lavage as described previously (Hu et al., 2015). At the termination of the experiment, selected organ tissues (kidney, lung, liver, spleen, stomach, and bladder) were harvested to determine whether viral infections could be detected in other unanticipated sites. The tissues were analyzed histologically for the presence of viral DNA and/or capsid protein production.

To investigate whether blood from actively infected mice could transmit the infection to naïve animals, blood was withdrawn from one infected male and one infected female mouse and transfused by tail vein injection to three naïve male and female mice respectively. The injection sites were treated topically with neutralizing antibody (MPV.A4) immediately post injection to neutralize any virions remaining at the site. The mice were maintained for up to six months post infection and tissues were collected for cellular, molecular, and histological analyses. Vaginal and anal lavages were conducted using 25 μl of sterile 0.9% NaCl introduced into the vaginal and anal canals with a disposable filter tip. The rinse was gently pipetted in and out of the canals and stored at −20°C before being processed for DNA extraction (Hu et al., 2015). For oral lavage, a swab (Puritan Purflock Ultra, Puritan Diagnostics LLC) soaked in 25 μl of sterile 0.9% NaCl was used (Cladel et al., 2017b). For DNA extraction, the DNeasy kit (QIAGEN) was used according to the instructions of the manufacturer. All DNA samples were eluted into 50 µl EB buffer (Hu et al., 2015).

### RNA isolation from rabbit tumors for quantitative PCR assays

Tumor tissues of NZW rabbits with CRPV infections induced by two different routes, a) local skin infection and b) ear IV injection, were harvested for QPCR analysis of host genes (Table 1). The tissues were homogenized in TriPure reagent (Roche). Total RNA was extracted according to the TriPure extraction protocol and treated with the TURBO DNA-free™ Kit (Ambion) to eliminate all traces of viral DNA. The integrity of RNA were evaluated by an Agilent DNA bioanalyzer and quantified by NanoDrop. Reverse Transcription (RT) was performed with the SuperScript II kit (Thermo Fisher Scientific). 1µg of total RNA from each tissue was used per reverse transcription reaction to synthesize single-stranded cDNA. cDNA samples were further analyzed using the TaqMan Universal PCR Master Mix (Thermo Fisher) by a StepOne Plus Real-Time PCR System (Applied Biosystems). To avoid interplate variability, differentially expressed gene expression analyses were performed using a single 96-well plate in triplicate. GAPDH was used as an internal control and analyzed in the same plate for each sample. Each threshold cycle (Ct) value of real-time quantitative PCR data from three repeats was individually normalized to GAPDH and analyzed by the 2-^ΔΔCt^ method (Xue et al., 2017). All primers and Taqman probes used were listed in Table 3.

### RNA-seq Analysis

Total RNA isolated using TriPure Reagent (Roche) and the RNeasy Mini kit with on-column DNase-treatment (Qiagen) was used for RNA-seq analysis. The sequencing libraries were constructed from Ribo-minus RNA using TruSeq Stranded Total RNA kit (Illumina RS-122-2201). The obtained libraries were then pooled and sequenced using Illumina TruSeq v4 chemistry, 125-bp paired-end, with 100 million reads depth. The HiSeq RT Analysis software (RTA v1.18.64) was used for base calling. The Illumina bclfastq v2.17 software was used to demultiplex and convert binary base calls and qualities to FASTQ format. The obtained reads were mapped first to the oryCun2.0 (*Oryctolagus cuniculus*) reference genome (http://www.ncbi.nlm.nih.gov/assembly/GCF_000003625.2/) to which had been added a contig containing the cottontail rabbit papillomavirus (CRPV) Hershey strain reference genome (GenBank Acc. No. JF303889.1) permutated at nt 7421. The viral read coverage along the CRPV reference genome was then visualized using the IGV software (http://software.broadinstitute.org/software/igv/). To determine the changes in host gene expression upon viral infection, the reads were remapped to the oryCun2.0 reference genome without the CRPV contig. RNA-seq NGS-datasets were processed using the CCBR Pipeliner utility (https://github.com/CCBR/Pipeliner). Briefly, reads were trimmed of low-quality bases and adapter sequences were removed using Trimmomatic v0.33 (Bolger et al., 2014). Mapping of reads to the oryCun2.0+CRPV reference genome was performed using STAR v2.5.2b in 2-pass mode (Dobin et al., 2013). Then, RSEM v1.3.0 was used to quantify gene-level expression, with counts normalized to library size as counts-per-million (Dobin et al., 2013). Finally, limma-voom v3.34.5 was used for quantile normalization and differential expression (Li and Dewey, 2011). The data discussed in this publication have been deposited in NCBI’s Gene Expression Omnibus (Phipson et al., 2016) and are accessible through GEO Series accession number GSE124211. Genes were considered to be attributed to CRPV infection if they were significantly (adjusted *p* ≤ 0.05) differentially expressed relative to control with absolute fold change relative to control ≥ 2.0. Genes with “unknown” gene symbols in the oryCun2.0 gene annotation dataset were quantified but excluded from further analysis in this manuscript. Expression data were visualized as heat maps using ClustVis (Metsalu and Vilo, 2015).

### Viral DNA copy number analysis

Linearized MmuPV1 genome DNA was used for standard curve determination by Probe qPCR analysis (Brilliant III Ultra-Fast QPCR Master Mix, Agilent). The primer pairs (5’-GGTTGCGTCGGAGAACATATAA-3’and 5’-CTAAAGCTAACCTGCCACATATC-3’) and the probe 5’-FAM-TGCCCTTTCA/ZEN/GTGGGTTGAGGACAG-3’-IBFQ-3’) that amplify the viral E2 region were used. The qPCR reactions were run in AriaMx program (Agilent). Each reaction consisted of 500 nM specific primer pairs and 250 nM double-labeled probes. PCR conditions were: initial denaturation at 95 °C for 10 min, then 40 cycles at 95°C for 10 min, followed by 40 cycles consisting of denaturation at 95°C for 15 s and hybridization of primers and the probe as well as DNA synthesis at 60 °C for 1 min. All samples were tested in at least duplicates. Viral titers were calculated according to the standard curve. Viral copy numbers in 2 μl of a 50 μl DNA lavage extract were converted into equivalent DNA load using the formula 1 ng viral DNA = 1.2 × 10^-8^ copy numbers (http://cels.uri.edu/gsc/cndna.html). In some cases we also calculated the difference in cycle time (Ct) between the 18S rRNA gene and viral DNA (ΔCt) (Hu et al., 2015). Fold change (2-^ΔΔ^Ct) demonstrates the relative viral DNA load in each sample as previously described (Cladel et al., 2017a).

### Antibody detection by ELISA

Rabbit and mouse sera were collected at the termination of the experiment. CRPV or MmuPV1 virus-like particles (VLPs) were used as the antigen for ELISA. Anti-CRPV monoclonal antibody (CRPV.1A) or anti-MmuPV1 monoclonal antibody (MPV.A4) was used as positive control and the sera of non-infected animals as negative control for the corresponding antigens. The ELISA was conducted as previously reported (Hu et al., 2014).

### In vitro neutralization assay

A rabbit cell line (RA2LT) generated in house was used for in vitro neutralization for serum collected from the CRPV infected rabbits (Hu et al., 2010). A mouse keratinocyte cell line (K38, a generous gift from Dr. Julia Reichelt, University of Newcastle, UK) was seeded at 1.5 ×10^-5^ cells per well in DMEM/Ham’s F-12, with 4.5 g/l D-Glucose, 50 uM CaCl2, with L-Glutamine and Na-Pyruvate (Cedarlane), in 10% FBS with calcium depleted at 32°C. One µl of viral extract from tail papillomas was incubated with various dilutions of mouse sera (1:50-1:100 dilution) in media for 1 hr at 37°C and added onto K38 cells incubated in 12-well plates at 32°C for 72 hours. The cells were harvested with TRIzol reagent (Thermo Fisher).

### CRPV E1^E4 detection by RT-qPCR

Total RNA was extracted from the infected cells, and infectivity was assessed by measuring viral E1^E4 transcripts with RT-qPCR (E1^E4-forward, 5’-CATTCGAGTC ACTGCTTCTGC-3’; E1^E4-reverse, 5’-GATGCAGGTTTGTCGTTCTCC-3’; E1^E4-probe, 5’-6-carboxyfluorescein (FAM)-TGGAAAACGATAAAGCTCCTCCTCAGCC-6-carboxytetramethylrhodamine (TAMRA)-3’ as previously described with a few modifications as follows: The Brilliant III RT-qPCR Master Mix (Agilent) was used for the RT-qPCR reactions. The following cycling conditions were applied: 50°C for 30 minutes (the reverse transcription), 95°C for 10 minutes, and 40 cycles of 94°C for 15 seconds and 60°C for 1 minute. At the end of each amplification cycle, three fluorescence readings were detected. Analysis of the amplification efficiencies was performed using the REST software (Cladel et al., 2017a).

#### Western blot analysis

Total protein from matching samples used in the RNA-seq study was isolated by homogenization in 1 × RIPA (Boston BioProducts) buffer supplemented by1 × complete protease inhibitors (Roche). The isolated total protein was analyzed by Western blot for the expression of endogenous protein using specific antibody against APOBEC2 (Sigma-Aldrich, cat. no. SAB2500083), S100A9 (Abnova, cat. no. PAB11470) and β-tubulin (Sigma-Aldrich, cat. no. T5201).

### Immunohistochemistry and in situ hybridization analyses of infected tissues

After termination of the experiment, the animals were euthanized, and tissues of interest were fixed in 10% buffered formalin and processed to formalin-fixed paraffin-embedded (FFPE) sections as previously described (Cladel et al., 2015). Hematoxylin and eosin (H&E) analysis, in situ hybridization (ISH) and immunohistochemistry (IHC) were conducted as described in previous studies (Cladel et al., 2015; Hu et al., 2015). For IHC, an in-house anti-MmuPV1 L1 monoclonal antibody (MPV.B9) was used on FFPE sections. For ISH, a biotin labeled 3913bp EcoRV/ BamHI sub genomic fragment of MmuPV1 was used as an in situ hybridization probe for the detection of MmuPV1 DNA in tissues (Cladel et al., 2015). Counterstaining for ISH was Nuclear Fast Red (American MasterTech, Inc.) and for IHC was hematoxylin (Thomas Scientific).

## ACKNOWLEDGEMENT

### Funding information

Research reported in this publication was supported by the NCI grant CA47622 to N.C., NIAID grant R21AI121822 to N.C. and J.H. and the Jake Gittlen Memorial Golf Tournament. This study was also supported by Intramural Research Program of NCI/NIH (1ZIASC010357 to Z.M.Z) and NCI/NIH contract (HHSN261200800001E). The content of this publication does not necessarily reflect the views or policies of the Department of Health and Human Services, nor does mention of trade names, commercial products, or organizations imply endorsement by the U.S. Government. This research was supported (in part) by the National Institutes of Health.

## Conflicts of interest

The authors declare that there are no conflicts of interest.

## Author Contributions

Conceptualization: ZZ, JH

Data curation: NMC, PJ, JL, XP, TKC, TJM, MC, KKB, SAB, VM, JH

Formal analysis: NMC, PJ, TKC, KKB, TJM, MC, VM, JH Funding acquisition: NDC, ZZ, JH

Investigation: NMC, JL, LRB, KKB, XP, TKC, DAS, SAB, RM, PJ, VM, RV, ZZ, NDC, JH

Methodology: NMC, LRB, KKB, PJ, JL, VM, SAB, TJM, MC, DAS, JH

Project administration: NMC, NDC, ZZ, JH Resources: NDC, ZZ, JH

Supervision: JH, ZZ, NDC

Validation: NMC, PJ, JL, DAS, TKC, LRB, TJM, MC, KKB, SAB, JH

Visualization: NMC, JL, LRB, TJM, KKB, TKC, DAS, JH

Writing – original draft: NMC, JH, ZZ

Writing – review & editing: NMC, JL, XP, TKC, VM, TJM, ZZ, NDC, JH

## Abbreviations

HPV: Human Papillomavirus
CRPV: The Cottontail Rabbit Papillomavirus
MmuPV1: The Mouse Papillomavirus
NZW: New Zealand White
IV: Intravenous
PBMCs: Peripheral Blood Mononuclear Cells
PCA: Principle Component Analysis
HCT: Hematopoietic Cell Transplantation

