## supplemental Fig 1, Table 1, 2 for "Papillomavirus can be transmitted through the blood and produce infections in blood recipients: Evidence from two animal models"

**Supplementary Figure 1.** HSD: Nu nude mice inoculated with anti-MmuPV1neutralizing antibody (MPV.A4) or a control monoclonal antibody (H11.B2 ) via i.p. at 24 hours before MmuPV1 infection at both cutaneous and mucosal sites. Viral infections can be detected at both cutaneous and mucosal by either visualization (B) or Q-PCR analysis (C) in the control group (H11.B2) but not in the test group (MPV.A4) (A, C) suggesting the neutralizing antibody can completely block viral infections in all the tested tissues.

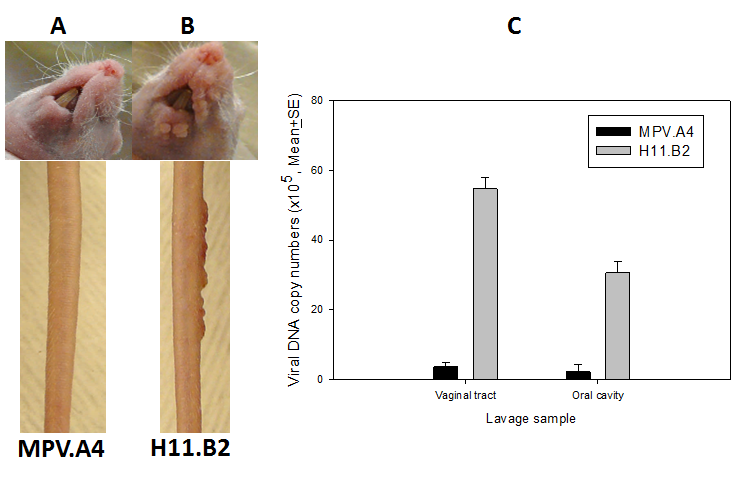

**Supplementary Table 1. Summary of CRPV virion titration at NZW domestic rabbit back sites**

| Viral titration | Viral genome equivalents | Papilloma appearance (tumor sites/infected skin sites) | |
| --- | --- | --- | --- |
|  |  | Without pre-wounding | With pre-wounding |
| 10^-1^ | 2.75×10_8_ | ND | 12/12 |
| 10^-2^ | 2.75×10_7_ | 5/5 | 33/33 |
| 10^-3^ | 2.75×10_6_ | 2/8 | 24/24* |
| 10^-4^ | 2.75×10_5_ | 0/16 | 28/36* |
| 10^-5^ | 2.75×10_4_ | 0/8 | 6/36 |
| 10^-6^ | 2.75×10_3_ | 0/8 | 1/20 |
| 10^-7^ | 2.75×10_2_ | ND | 0/12 |

*P<0.01 with pre-wounding vs. without pre-wounding, Fisher’s exact test

**Supplementary Table 2. Summary of CRPV DNA titration at NZW domestic rabbit back sites**

| Viral DNA (µg) | Viral genome equivalents | Papilloma appearance (tumor sites/infected skin sites) | | |
| --- | --- | --- | --- | --- |
|  |  | Without pre-wounding | | With pre-wounding |
| 20 | 2.6×10^12^ | ND | 28/28 | |
| 10 | 1.3×10_12_ | 3/5 | 4/5 | |
| 5 | 6.5×10_11_ | 5/8 | 11/11 | |
| 1 | 1.3×10_11_ | 4/5 | 11/11 | |
| 0.2 | 2.6×10_10_ | 2/5 | 11/11* | |
| 0.1 | 1.3×10_10_ | 2/5 | 3/3 | |
| 0.04 | 5.2×10_9_ | 3/5 | 9/11 | |
| 0.02 | 2.6×10_9_ | 0/3 | 1/3 | |
| 0.01 | 1.3×10_9_ | 1/3 | 1/3 | |

*P<0.05 with pre-wounding vs. without pre-wounding, Fisher’s exact test

**Supplementary Table 3** The dysregulated genes in RNAseq analysis.
